# Resistance of SARS-CoV-2 Omicron Subvariant BA.4.6 to Antibody Neutralization

**DOI:** 10.1101/2022.09.05.506628

**Authors:** Qian Wang, Zhiteng Li, Jerren Ho, Yicheng Guo, Andre Yanchen Yeh, Michael Liu, Maple Wang, Jian Yu, Zizhang Sheng, Yaoxing Huang, Lihong Liu, David D. Ho

## Abstract

SARS-CoV-2 Omicron subvariants BA.4.6, BA.4.7, and BA.5.9 have recently emerged, and BA.4.6 appears to be expanding even in the presence of BA.5 that is globally dominant. Compared to BA.5, these new subvariants harbor a mutation at R346 residue in the spike glycoprotein, raising concerns for further antibody evasion. We compared the viral receptor binding affinity of the new Omicron subvariants with BA.5 by surface plasmon resonance. We also performed VSV-based pseudovirus neutralization assays to evaluate their antigenic properties using sera from individuals who received three doses of a COVID-19 mRNA vaccine (boosted) and patients with BA.1 or BA.2 breakthrough infection, as well as using a panel of 23 monoclonal antibodies (mAbs). Compared to the BA.5 subvariant, BA.4.6, BA.4.7, and BA.5.9 showed similar binding affinities to hACE2 and exhibited similar resistance profiles to boosted and BA.1 breakthrough sera, but BA.4.6 was slightly but significantly more resistant than BA.5 to BA.2 breakthrough sera. Moreover, BA.4.6, BA.4.7, and BA.5.9 showed heightened resistance over to a class of mAbs due to R346T/S/I mutation. Notably, the authorized combination of tixagevimab and cilgavimab completely lost neutralizing activity against these three subvariants. The loss of activity of tixagevimab and cilgavimab against BA.4.6 leaves us with bebtelovimab as the only therapeutic mAb that has retained potent activity against all circulating forms of SARS-CoV-2. As the virus continues to evolve, our arsenal of authorized mAbs may soon be depleted, thereby jeopardizing the wellbeing of millions of immunocompromised persons who cannot robustly respond to COVID-19 vaccines.

## Main text

Severe acute respiratory syndrome coronavirus 2 (SARS-CoV-2), the causative agent of the coronavirus disease 2019 (COVID-19) pandemic, continues to evolve. An Omicron subvariant known as BA.4.6 has recently emerged, and it appears to be expanding even in the presence of BA.5, another subvariant that has been globally dominant in recent months^1,2^ (Fig. S1A). Compared to BA.4 or BA5, BA.4.6 contains two additional mutations, R346T and N658S, in the spike protein (Fig. S1B). Two other nascent Omicron subvariants with similar spike mutations, BA.4.7 with R346S and BA.5.9 with R346I, have also been detected, albeit at extremely low frequencies (Figs. S1A and S1B). That these three new subvariants all have mutations at the R346 residue raises concerns for further antibody evasion, because R346K in a prior subvariant (BA.1.1) impaired the potency of several therapeutic monoclonal antibodies (mAbs)^3,4^.

We first examined whether the transmission advantage of BA.4.6 could be due to a higher affinity for the viral receptor. Affinity measurements were made for the binding of purified spike trimers of D614G, BA.2, BA.4/5, BA.4.6, BA.4.7, and BA.5.9 to dimeric human ACE2 (hACE2) by surface plasmon resonance (Figs. 1A and S2). All the spike proteins from BA.4/5 sublineages, as well as those of BA.4/5 carrying point mutations of R346T, R346S and N658S, showed similar binding affinities to hACE2, with KD values ranging from 0.39 nM to 0.49 nM. Therefore, the expansion of BA.4.6 cannot be explained by a higher affinity for hACE2.

**Figure 1.**
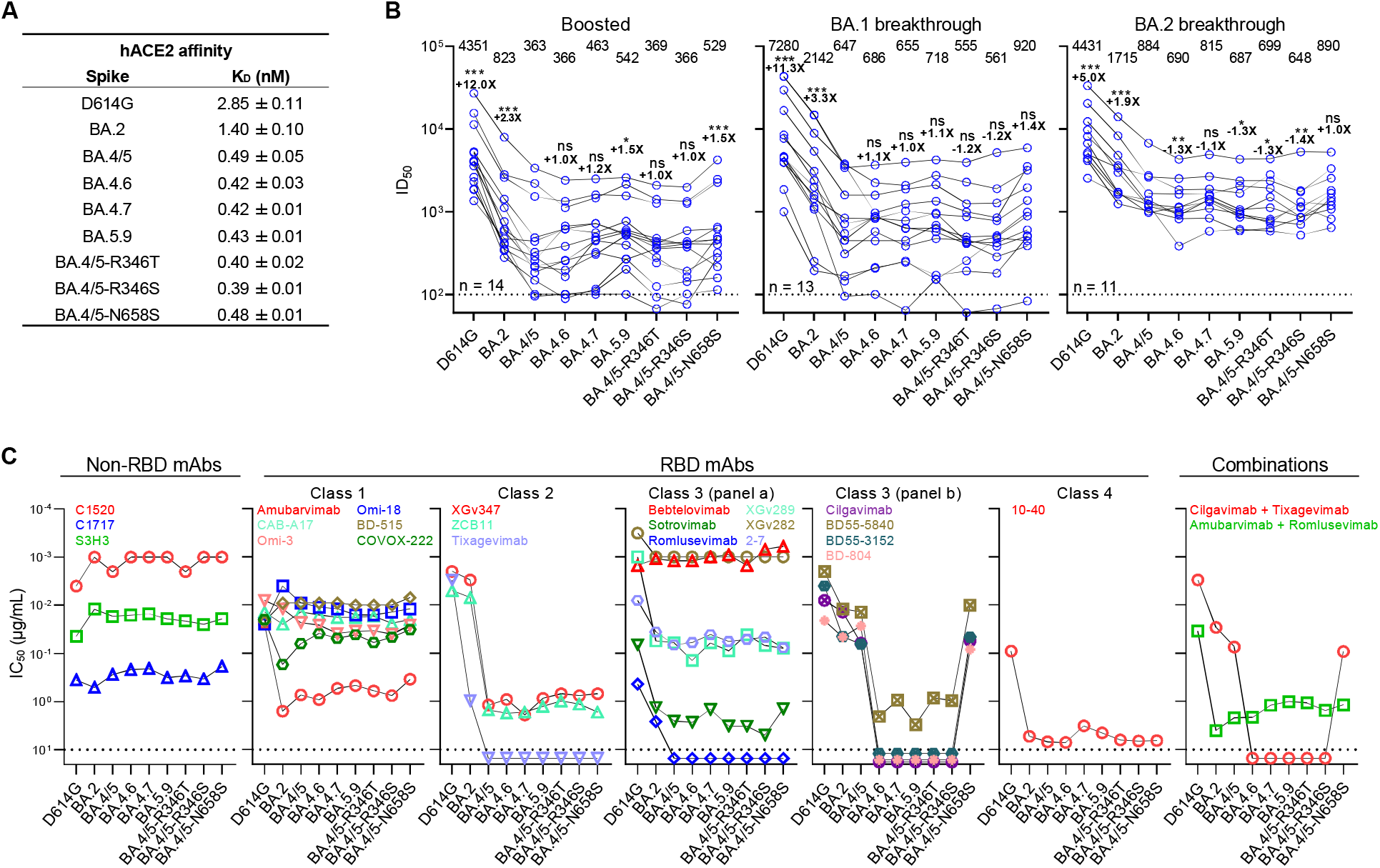
Receptor binding affinity and antibody neutralization profiles of new Omicron subvariants. Panel **A** shows the ACE2 binding affinity of the spikes of BA.4.6, BA.4.7, and BA.5.9, as well as the individual mutations found in BA.4.6 and BA.4.7 in the background of BA.4/5. D614G, BA.2, and BA.4/5 spikes were included for comparison. Data are shown as K_D_ values. Panel **B** shows the neutralization ID_50_ titers of the serum samples from the “boosted”, “BA.1 breakthrough”, and “BA.2 breakthrough” cohorts. Values above the symbols denote the geometric mean ID_50_ values and values on the lower left indicate the sample size (n). The limit of detection is 100 (dotted line). Comparisons were made against BA.4/5 by two-tailed Wilcoxon matched-pairs signed-rank tests. *, *P* < 0.05; **, *P* < 0.01; ***, *P* < 0.001; and ns, not significant. Panel **C** shows the neutralization by mAbs of D614G, Omicron subvariants, and point mutants in the background of BA.4/5. Values above the maximum antibody concentration tested, 10□μg/ml. (dotted line), are arbitrarily plotted to allow for visualization of each sample. Preclinical mAbs are denoted by their laboratory designations, and clinical mAbs are denoted by their generic names. The combination of cilgavimab and tixagevimab is marketed as Evusheld.

To assess the antibody evasion properties of BA.4.6, BA.4.7, and BA.5.9, we evaluated the sensitivity of their corresponding pseudoviruses (see Methods in the Supplement) to neutralization by serum samples from healthy individuals who had received three doses of a COVID-19 mRNA vaccine (boosted) and patients with either BA.1 or BA.2 breakthrough infection after mRNA vaccination (Table S1). The results are shown in Fig. 1B. The ID_50_ (50% inhibitory dose) titers of the boosted samples against BA.4.6, BA.4.7, and BA.5.9 were similar to that against BA.4/5, with no more than 1.5-fold deviation in the geometric mean values. Likewise, the individual mutations R346T, R346S, and N658S in the background of BA.4/5 had little impact on the neutralization profiles. A similar trend was also observed in serum neutralization for BA.1 breakthrough cases. Against sera from the BA.2 breakthrough cohort, BA.4.6 was slightly (1.3-fold) but significantly (P <0.01) more resistant than BA.5, although it remains unclear if this small difference could explain the recent expansion of BA.4.6 worldwide.

To further characterize the antigenic properties of BA.4.6, along with BA.4.7 and BA.5.9, we measured the sensitivity of each subvariant pseudovirus to neutralization by a panel of 23 mAbs that retained potency against earlier Omicron subvariants, including ones that target different epitope clusters (classes 1, 2, 3, and 4) of the receptor-binding domain (RBD) of the viral spike and others that target non-RBD epitopes (Figs. 1C and S3). In general, the neutralization profiles of BA.4.6, BA.4.7, and BA.5.9 did not differ much from that of BA.4/5. The only exceptions were mAbs in RBD class 3 (panel b), which showed marked reduction in their neutralization potency against the new subvariants. This loss of neutralizing activity was due to mutation R346T or R346S but not due to N658S. Structural analyses revealed that R346T/S/I mutations eliminated or weakened hydrogen bonds and/or salt bridges between R346 and certain RBD class 3 mAbs (Fig. S4), explaining why these mutations led to substantial neutralization resistance. These findings suggest that BA.4.6, BA.4.7 and BA.5.9 likely emerged under the selective pressure of RBD class 3 antibodies in infected individuals.

Importantly, several mAbs in clinical use were included in the neutralization assays against the new Omicron subvariants as well (Fig. 1C). The combination of cilgavimab and tixagevimab, which had received emergency use authorization for the prevention of COVID-19^5^, could not neutralize BA.4.6, nor BA.4.7 and BA.5.9. The loss of this antibody combination against BA.4.6 leaves us with bebtelovimab as the only therapeutic monoclonal antibody that has retained potent activity against all circulating forms of SARS-CoV-2. As the pandemic rages on and as the virus continues to evolve, our arsenal of authorized monoclonal antibodies may soon be depleted, thereby jeopardizing the wellbeing of millions of immunocompromised persons who cannot robustly respond to COVID-19 vaccines.

## Notes

### Competing Interest Statement

J.Y., Y.H., L.L., and D.D.H. are inventors on patent applications (WO2021236998) or provisional patent applications (63/271,627) filed by Columbia University for a number of SARS-CoV-2 neutralizing antibodies described in this manuscript. Both sets of applications are under review. D.D.H. is a co-founder of TaiMed Biologics and RenBio, consultant to WuXi Biologics and Brii Biosciences, and board director for Vicarious Surgical.

## Reference

1. Shu Y, McCauley J. GISAID: Global initiative on sharing all influenza data - from vision to reality. Euro Surveill 2017;22.

2. COVID Data Tracker. Atlanta, GA: US Department of Health and Human Services, CDC, 2022. (Accessed 2022, August 31, at https://covid.cdc.gov/covid-data-tracker.)

3. Liu L, Iketani S, Guo Y, et al. Striking antibody evasion manifested by the Omicron variant of SARS-CoV-2. Nature 2022;602:676–81.

4. Wang Q, Guo Y, Iketani S, et al. Antibody evasion by SARS-CoV-2 Omicron subvariants BA.2.12.1, BA.4, & BA.5. Nature 2022.

5. FDA authorizes revisions to Evusheld dosing. U.S. Food & Drug Administration, 2022. (Accessed 2022, September 1, at https://www.fda.gov/drugs/drug-safety-and-availability/fda-authorizes-revisions-evusheld-dosing.)

