## Supplemental figures for "Resistance of SARS-CoV-2 Omicron Subvariant BA.4.6 to Antibody Neutralization"

### Supplementary Methods

*Patients and vaccinees*

Sera from individuals who were vaccinated with three doses of the mRNA-1273 or BNT162b2 vaccine were collected at Columbia University Irving Medical Center (referred to as “boosted” in the text). Sera from individuals who received mRNA vaccinations and were subsequently infected by Omicron subvariant BA.1 or BA.2 were collected at Columbia University Irving Medical Center from December 2021 to May 2022 (referred to as “BA.1 or BA.2 breakthrough” in the text). All samples were examined by anti-nucleoprotein (NP) ELISA to confirm status of prior SARS-CoV-2 infection, and the sequencing was conducted to determine the viral genotype. All subjects provided written informed consent, and all serum collections were performed under protocols reviewed and approved by the Institutional Review Board of Columbia University. Clinical information on different study cohorts is provided in Table S1.

*Cell lines*

HEK293T cells (CRL-3216) and Vero-E6 cells (CRL-1586) were obtained from the ATCC and were maintained in Dulbecco modified Eagle medium (DMEM) with 10% fetal bovine serum (FBS) and 1% penicillin-streptomycin in an atmosphere of 5% CO_2_ at 37°C. Expi293 cells were obtained from Thermo Fisher Scientific (A14527) and were maintained in Expi293^TM^ Expression Medium supplemented with 0.5% penicillin-streptomycin (Thermo Fisher Scientific, Cambridge, MA) at 37°C, 8% CO_2_, and 125 rpm.

*Monoclonal antibodies*

Antibodies were expressed in-house as previously described^1^. Genes of the heavy chain variable (VH) and light chain variable (VL) for each antibody were synthesized (GenScript), cloned into an expression vector (pCMV3 or gWiz), transfected into Expi293 cells using polyethylenimine (PEI), and purified from the supernatants with affinity purification using rProtein A Sepharose (GE) on the fourth day after transfection. Cilgavimab and tixagevimab were obtained from Regeneron Pharmaceuticals.

*Construction of SARS-CoV-2 spike plasmids*

Plasmids containing spikes of D614G, BA.2, and BA.4/5 were previously constructed^2-4^. Expression constructs of the BA.4.6 spike, along with the individual mutations found in BA.4.6, were produced with the QuikChange II XL site-directed mutagenesis kit according to the manufacturer’s instructions (Agilent). To express stabilized soluble spike trimer proteins, 2P substitutions (K986P and V987P in WA1) and a “GSAS” substitution at the furin cleavage site (682-685aa in WA1) were introduced into the spike expression constructs as previously described^5^. The ectodomain (1-1208aa in WA1) of the spike was then fused with a C-terminal 8x His-tag and cloned into the paH vector. All constructs were confirmed by Sanger sequencing prior to experiments.

*Expression and purification of SARS-CoV-2 stabilized spike trimers and human ACE2*

paH-spike or pcDNA3-sACE2-WT(732)-IgG1 (Addgene plasmid #154104) plasmid was transfected into Expi293 cells using PEI as the transfection reagents, and the supernatants were harvested five days after. To purify the spike proteins, Tris-HCl (PH=8.0) buffer was added at a final concentration of 20mM, and then the solution flowed through the Excel resin (Cytiva) according to the manufacturer’s instructions. For purification of the human ACE2, Protein A Sepharose (Cytiva) was used following the manufacturer’s instructions. All proteins were confirmed by SDS-PAGE, with a purity of approximately 95% before experimental use.

*Surface plasmon resonance (SPR)*

SPR experiments were performed with a Biacore T200 system equipped with CM5 chips (Cytiva) at ambient temperature (25℃). The anti-His antibodies were immobilized on the CM5 chip by the His Capture Kit (Cytiva) to reach around 10000 RU. The spike protein was captured on the chip through the C-terminal His-tag, and human ACE2-Fc proteins then flowed through the chip surface at a gradient concentration in HBS-EP+ buffer (Cytiva). The single cycle binding kinetics was analyzed by the Evaluation Software using the 1:1 binding model.

*Pseudovirus production*

Pseudotyped SARS-CoV-2 were generated in the background of vesicular stomatitis virus (VSV), whose native VSV glycoprotein was replaced with those of SARS-CoV-2 variants as previously described^1^. HEK293T cells were transfected with plasmids containing the appropriate spike using 1 mg mL^-1^ of PEI. The transfected HEK293T cells were cultured under 5% CO_2_ at 37 °C for 24 hours, and then they were infected with VSV-G pseudotyped ΔG-luciferase (G*ΔG-luciferase, Kerafast). Two hours after, the infected HEK293T cells were washed three times before being cultured in fresh medium for another 24 hours. The supernatants were then collected, centrifuged to remove precipitates, and aliquoted for storage at -80 °C until the next use.

*Pseudovirus neutralization assay*

All pseudoviruses were titrated to equilibrate the viral input before each round of neutralization assay. Heat-inactivated sera or antibodies were serially diluted at five-fold in media in triplicate in 96-well plates, starting at 1:100 dilution for sera and 10 µg mL^−1^ for antibodies. Pseudoviruses were added and the virus–sample mixture was incubated at 37 °C for 1 hour. Control wells that only contained the virus were included on all plates. Vero-E6 cells were then added at a density of 3 × 10^4^ cells per well and the plates were incubated at 37 °C for 10 hours. Cells were lysed and luciferase activity was measured using the Luciferase Assay System (Promega) and SoftMax Pro v.7.0.2 (Molecular Devices) according to instructions from the two manufacturers. ID_50_ and IC_50_ values were obtained by fitting a nonlinear five-parameter dose-response curve to the data in GraphPad Prism v.9.2.

*Antibody footprint analysis and structural modeling of RBD mutations*

The structures of antibody-spike complexes were obtained from Protein Data Bank (6WPS for sotrovimab, 7LSS for 2-7, 7MMO for bebtelovimab, 7WEF for XGv289, 7WLC for XGv282, 7X6A for BD55-5840, 7WR8 for BD55-3152, and 7EYA for BD-804) for modeling. The interface residues are generated by using the script from PyMOLwiki. The boundaries of all epitope residues are defined as the antibody footprint and then optimized in Adobe Photoshop. The interaction residues are obtained by CCP4. PyMOL v.2.3.2 was used to perform mutagenesis, to identify hydrogen bonds as well as salt bridge between RBD and antibodies, and to generate structural plots (Schrödinger, LLC).

*Quantification and statistical analysis*

Serum neutralization ID_50_ values and antibody neutralization IC_50_ values were obtained from a five-parameter dose-response curve in GraphPad Prism v.9.2. Statistical significance was evaluated by two-tailed Wilcoxon matched-pairs signed-rank tests using GraphPad Prism v.9.2. Levels of significance are denoted as follows: ns, not significant; *, *P* < 0.05; **, *P* < 0.01; and ***, *P* < 0.001.

### Acknowledgements

This study was supported by funding from the Gates Foundation, JPB Foundation, Andrew and Peggy Cherng, Samuel Yin, Carol Ludwig, David and Roger Wu, Regeneron Pharmaceuticals, and the NIH SARS-CoV-2 Assessment of Viral Evolution (SAVE) Program. We are grateful to Michael T. Yin, Magdalena E. Sobieszczyk, Jennifer Y. Chang, Jayesh G. Shah, and David S. Perlin for providing serum samples from COVID-19 patients.

### Author Contributions

D.D.H. and L.L. conceived this project. Q.W., A.Y.Y., and L.L. constructed the spike expression plasmids and produced pseudoviruses. Q.W. and L.L. conducted pseudovirus neutralization experiments. Q.W., J.H., and L.L. purified SARS-CoV-2 spike and ACE2 proteins. Z.L. performed SPR. Z.L., Y.G., and Z.S. conducted bioinformatic analyses. Q.W. managed the project. J.H., M.L., J.Y., and M.L. produced antibodies. D.D.H. and L.L. directed and supervised the project. Q.W., L.L., and D.D.H. analyzed the results and wrote the manuscript.

### Declaration of interests

J.Y., Y.H., L.L., and D.D.H. are inventors on patent applications (WO2021236998) or provisional patent applications (63/271,627) filed by Columbia University for a number of SARS-CoV-2 neutralizing antibodies described in this manuscript. Both sets of applications are under review. D.D.H. is a co-founder of TaiMed Biologics and RenBio, consultant to WuXi Biologics and Brii Biosciences, and board director for Vicarious Surgical.

### Table S1. Demographics of clinical cohorts in this study.

| Sample ID | Vaccine type and infected strain | Days post-vaccination or *infection  (after last exposure) | Documented COVID-19 | Age | Gender |
| --- | --- | --- | --- | --- | --- |
| Boosted | | | | | |
| Q1 | mRNA-1273/mRNA-1273/mRNA-1273 | 29 | No | 66 | Female |
| Q2 | BNT162b2/BNT162b2/BNT162b2 | 30 | No | 68 | Male |
| Q3 | BNT162b2/BNT162b2/BNT162b2 | 14 | No | 64 | Female |
| Q4 | BNT162b2/BNT162b2/BNT162b2 | 34 | No | 55 | Male |
| Q5 | BNT162b2/BNT162b2/BNT162b2 | 34 | No | 45 | Male |
| Q6 | BNT162b2/BNT162b2/BNT162b2 | 15 | No | 50 | Female |
| Q7 | BNT162b2/BNT162b2/BNT162b2 | 15 | No | 48 | Female |
| Q8 | BNT162b2/BNT162b2/BNT162b2 | 29 | No | 71 | Male |
| Q9 | BNT162b2/BNT162b2/BNT162b2 | 90 | No | 59 | Male |
| Q10 | BNT162b2/BNT162b2/BNT162b2 | 33 | No | 45 | Male |
| Q11 | BNT162b2/BNT162b2/BNT162b2 | 87 | No | 66 | Female |
| Q12 | BNT162b2/BNT162b2/BNT162b2 | 84 | No | 26 | Male |
| Q13 | mRNA-1273/mRNA-1273/mRNA-1273 | 23 | No | 28 | Female |
| Q15 | BNT162b2/BNT162b2/mRNA-1273 | 32 | No | 39 | Male |
| BA.1 breakthrough | | | | | |
| Q24 | BNT162b2/BNT162b2/BA.1 | *14 | Yes | Unknown | Unknown |
| Q25 | BNT162b2/BNT162b2/BA.1 | *14 | Yes | Unknown | Unknown |
| Q26 | mRNA-1273/mRNA-1273/BA.1 | *35 | Yes | Unknown | Unknown |
| Q27 | BNT162b2/BNT162b2/BNT162b2/BA.1 | *135 | Yes | 78 | Male |
| Q28 | BNT162b2/BNT162b2/BNT162b2/BA.1 | *14 | Yes | Unknown | Unknown |
| Q29 | BNT162b2/BNT162b2/BNT162b2/BA.1 | *14 | Yes | Unknown | Unknown |
| Q30 | BNT162b2/BNT162b2/BNT162b2/BA.1 | *14 | Yes | Unknown | Unknown |
| Q31 | BNT162b2/BNT162b2/BNT162b2/BA.1 | *41 | Yes | 48 | Male |
| Q32 | BNT162b2/BNT162b2/BNT162b2/BA.1 | *26 | Yes | 38 | Female |
| Q33 | BNT162b2/BNT162b2/B.1.617.2/BNT162b2/BA.1 | *19 | Yes | 35 | Female |
| Q34 | BNT162b2/BNT162b2/mRNA-1273/mRNA-1273/BA.1 | *67 | Yes | 40 | Male |
| Q41 | WA1/BNT162b2/BA.1 | *21 | Yes | 52 | Male |
| Q42 | WA1/BNT162b2/BA.1 | *44 | Yes | 37 | Intersex |
| BA.2 breakthrough | | | | | |
| Q35 | BNT162b2/BNT162b2/BA.2 | *14 | Yes | 50 | Female |
| Q36 | BNT162b2/BNT162b2/BNT162b2/Ad26.COV2.S/BA.2 | *22 | Yes | 69 | Male |
| Q50 | mRNA-1273/mRNA-1273/mRNA-1273/BA.2 | *14 | Yes | 34 | Male |
| Q51 | BNT162b2/BNT162b2/mRNA-1273/BA.2 | *19 | Yes | 33 | Female |
| Q52 | BNT162b2/BNT162b2/mRNA-1273/BA.2 | *18 | Yes | 29 | Female |
| Q53 | BNT162b2/BNT162b2/BNT162b2/BA.2 | *25 | Yes | 34 | Male |
| Q54 | BNT162b2/BNT162b2/BNT162b2/BA.2 | *36 | Yes | 37 | Female |
| Q55 | BNT162b2/BNT162b2/mRNA-1273/BA.2 | *18 | Yes | 41 | Female |
| Q56 | mRNA-1273/mRNA-1273/mRNA-1273/BA.2 | *21 | Yes | 36 | Female |
| Q57 | BNT162b2/BNT162b2/mRNA-1273/BA.2 | *32 | Yes | 28 | Male |
| Q58 | BNT162b2/BNT162b2/mRNA-1273/BA.2 | *23 | Yes | 33 | Female |

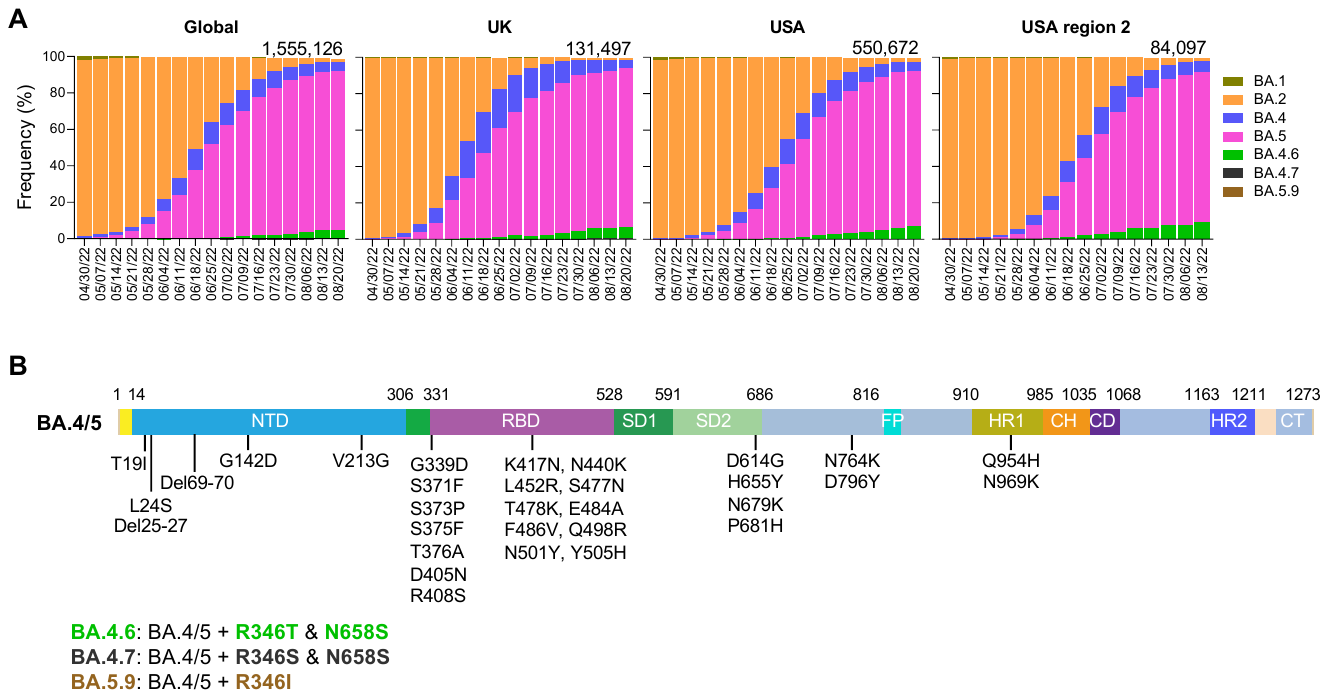

### Figure S1. Prevalence of Omicron subvariants and mutations found in BA.4.6 spike.

A. Frequencies of Omicron subvariants deposited in GISAID. The value in the upper right corner of each box shows the cumulative number of sequences for all circulating viruses in the denoted time period. USA region 2 includes New Jersey, New York, Puerto Rico, and Virgin Islands.
B. Spike mutations found in BA.4.6 relative to BA.4/5. NTD, N-terminal domain; RBD, receptor-binding domain; SD1 and SD2, subdomains 1 and 2. FP, fusion peptide; HR1 and HR2, heptad repeat region 1 and 2; CH, central helical region; CD, connector domain; TM, transmembrane region; CT, cytoplasmic tail. The amino acid changes in the BA.4/5 spike relative to the D614G are shown in the diagrams.

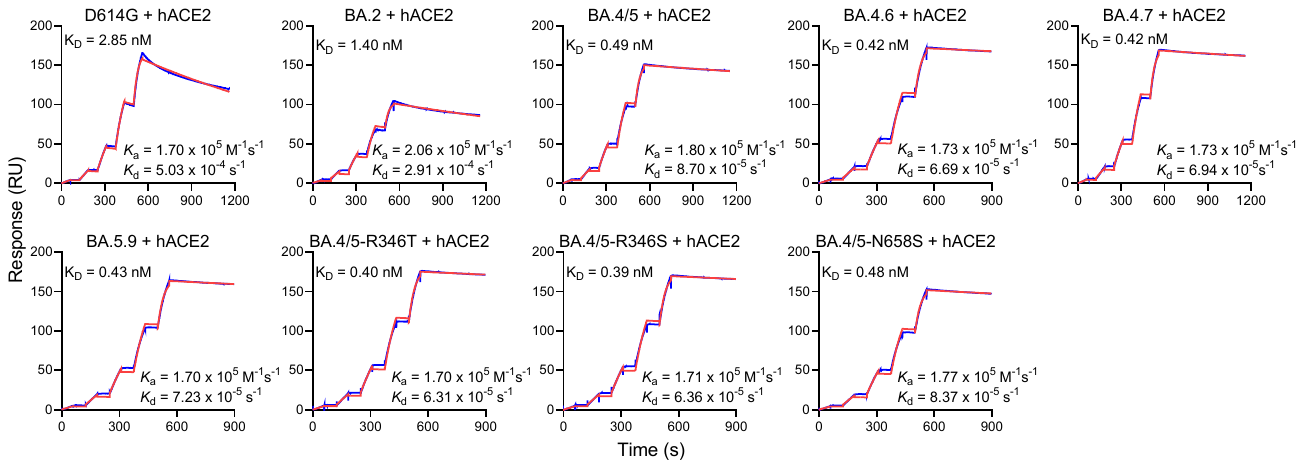

### Figure S2. Binding affinities of Omicron subvariant spike trimer proteins to hACE2 as measured by SPR.

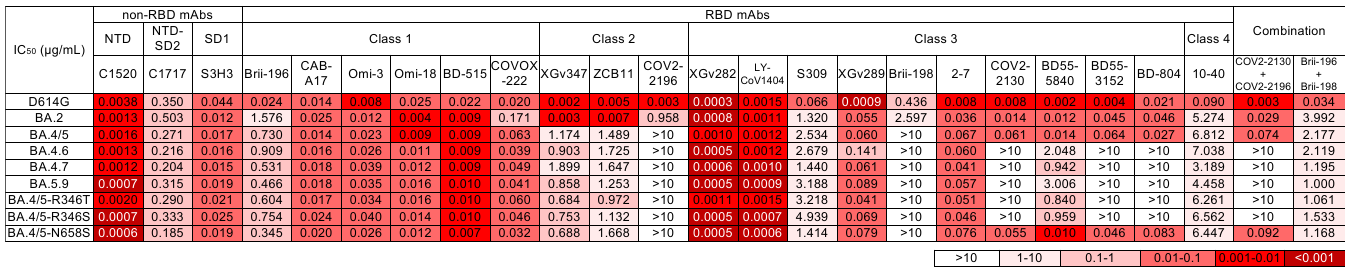

### Figure S3. Neutralization IC_50_ titers for indicated pseudoviruses by mAbs. The brand names of therapeutic neutralizing antibodies in the table are: Brii-196 (amubarvimab), COV2-2196 (tixagevimab), LY-CoV1404 (bebtelovimab), S309 (sotrovimab), Brii-198 (romlusevimab), COV2-2130 (cilgavimab). Antibody combinations: Evusheld consists of tixagevimab co-packaged with cilgavimab, and the Brii cocktail combination consists of amubarvimab and romlusevimab. Background colors indicate neutralization levels.

### Figure S4. Modeling of how R346T affects RBD class 3 mAb neutralization.

A. Footprints of class 3 neutralizing mAbs on the RBD.
B. Structural analysis for how R346T affects RBD class 3 mAbs binding. The green dashed lines denote the hydrogen bond and the salt bridge.

### Supplementary References

1.	Liu L, Wang P, Nair MS, et al. Potent neutralizing antibodies against multiple epitopes on SARS-CoV-2 spike. Nature 2020;584:450-6.

2.	Liu L, Iketani S, Guo Y, et al. Striking antibody evasion manifested by the Omicron variant of SARS-CoV-2. Nature 2022;602:676-81.

3.	Iketani S, Liu L, Guo Y, et al. Antibody evasion properties of SARS-CoV-2 Omicron sublineages. Nature 2022;604:553-6.

4.	Wang Q, Guo Y, Iketani S, et al. Antibody evasion by SARS-CoV-2 Omicron subvariants BA.2.12.1, BA.4, & BA.5. Nature 2022.

5.	Wrapp D, Wang N, Corbett KS, et al. Cryo-EM structure of the 2019-nCoV spike in the prefusion conformation. Science 2020;367:1260-3.
